# Prostaglandin F_2_α-induced TGFβ-PMEPA1 pathway is a critical mediator of epithelial plasticity and ovarian carcinoma progression

**DOI:** 10.1101/418582

**Authors:** Alba Jiménez-Segovia, Alba Mota, Alejandro Rojo-Sebastián, Beatriz Barrocal, Angela Rynne-Vidal, María-Laura García-Bermejo, Raquel Gómez-Bris, Lukas J.A.C. Hawinkels, Pilar Sandoval, Ramon Garcia-Escudero, Manuel López-Cabrera, Gema Moreno-Bueno, Manuel Fresno, Konstantinos Stamatakis

## Abstract

Prostaglandin (PG) F_2α_ has been scarcely studied in cancer. We have identified a new role for PGF_2α_ in ovarian cancer, stimulating the production of TGFβ and the consequent induction of PMEPA1. We show that this induction increases cell plasticity and proliferation, enhancing tumor growth through PMEPA1. Thus, PMEPA1 overexpression in ovarian carcinoma cells, significantly increased cell proliferation rates, whereas PMEPA1 silencing decreased proliferation. In addition, PMEPA1 overexpression buffered TGFβ signaling, via reduction of SMAD-dependent signaling. PMEPA1 overexpressing cells acquired an epithelial morphology, associated to higher E-cadherin expression levels while β-catenin nuclear translocation was inhibited. Interestingly, in mouse xenografts, PMEPA1 overexpressing ovarian cells had a clear survival and proliferative advantage, resulting in higher metastatic capacity, while PMEPA1 silencing had the opposite effect. Furthermore, high PMEPA1 expression in a cohort of advanced ovarian cancer patients was observed, correlating with E-cadherin expression. Most importantly, high PMEPA1 mRNA levels were associated with lower patient survival.

**Significance:** This work identifies Prostaglandin F_2α_ as an inducer, through NFAT, of TGFβ and together with this up-regulate PMEPA1, which is identified both as a drug target and as a prognostic biomarker in ovarian tumors, potentially useful in clinical practice.

## Introduction

The cyclooxygenases (COX) are enzymes that catalyze the rate limiting step in the biosynthesis of prostaglandins (PGs), which have also been implicated in various disease states including cancer (1,2). Clinical trials have shown that COX inhibitors (NSAIDs) reduce the risk of developing cancer and associated death (3). In this regard, higher COX2 expression and PG levels have been detected in different tumors, which could explain the NSAIDs beneficial effects in cancer prevention and treatment (3). Prostaglandin F_2α_ (PGF_2α_) has been scarcely studied on cancer although it has been detected in several tumor types and cancer patient body fluids (4,5) and only recently has been mechanistically associated with cancer progression (6). PGF_2α_ is produced through the combined action of cyclooxygenases (COX) and different isomerases, AKR1B1 and AKR1C3 among others, with scarce reports implicating them in cancer (7). PGF_2α_ binds and signals through the prostaglandin F receptor (PTGFR), a G protein coupled receptor, coupling mainly to Gq, thus leading to an increase in intracellular Ca^2+^ levels (8). Consequently, PGF_2α_ binding to FP will cause Ca^2+^/calcineurin activation of the NFAT transcription factors, implicated in a growing variety of physiological and pathological functions, including cancer (9).

On the other hand, previous studies have shown there are cell and stage specific increases of COX, prostaglandin synthases and receptors in epithelial ovarian cancer (EOC), supporting the hypothesis that PG synthesis and signaling are of importance for malignant transformation and progression in EOC (10,11). EOC, which comprises 90% of all ovarian malignancies, is the leading cause of death from gynecological cancer, in developed countries (12). Moreover, the majority of patients are diagnosed in advanced cancer stages and the average 5-year survival rate is low (13). Increased contents of prostaglandins in ovarian tumors have been described in previous studies, possibly regulating cell proliferation and apoptosis in this specific tumor context (11), being PGE_2_ the most extensively studied in this context (10). However, we have found another PG with an emerging role in colon cancer progression (6) with a close relationship with the female reproductive system (14).

PMEPA1 (also known as *TMEPA1*, transmembrane prostate androgen induced 1) gene was first identified as a highly androgen-inducible gene with abundant expression in prostatic epithelial cells, which eventually was described as a direct transcriptional target of the androgen receptor (AR) (15) but also being upregulated in renal cell carcinoma (16). *PMEPA1* expression, has been found in primary and metastatic pancreatic, endometrial, ovarian, colorectal, breast, lung and prostate tumors (17-20) although its role has been scarcely explored.

*PMEPA1* is known to be induced by transforming growth factor-β (TGF-β) (18). The TGF-β cytokine and the related signaling pathway are key players in mammalian health and disease, with an extensively studied, yet controversial role in cancer (21). Moreover, TGF-β has been shown to control ovarian cancer cell proliferation (22). PMEPA1 downregulates TGF-β signaling by competing with SARA (SMAD anchor for receptor activation) for R-SMAD binding to sequester R-SMAD phosphorylation and promoting lysosomal degradation of TGF-β receptor(23). PMEPA1, through a negative feedback loop, is described as the responsible to switch TGFβ from a tumor suppressor to a tumor promoter in breast cancer (20). In addition, TGF-β-dependent growth of aggressive breast cancer has been suggested to depend on increased expression of *PMEPA1* gene (19).

*PMEPA1* has been also described as a Wnt/β-catenin cascade target gene (24,25), which aberrant activation could contribute to neoplasia. In the case of endometrioid ovarian carcinomas, this stabilization is due to mutations identified in the *CTNNB1* gene (26), while for the rest, this pathway remains unaltered (27). β-catenin also binds with the intracellular domain of epithelial-cadherin (E-Cadherin) to form a complex which subsequently connects to the actin cytoskeleton, mediates intercellular adhesion, and regulates tumor cell invasion and metastasis (28).

Several oncogenic pathways (i.e., Wnt/β-catenin) increase cell plasticity (29). TGF-β is a potent regulator of cell plasticity through the induction of epithelial to mesenchymal transition (EMT) (30), directly activating the expression of transcription factors such as SNAI1/2, Twist and ZEB1/2, which are the key regulators of the EMT program. Related to this, SMADs (SMAD2 and SMAD3 particularly) have been proposed as critical mediators in EMT (31,32). In view of this information, and since PMEPA1 is described as SMAD2/3 interacting protein, affecting TGF-β target genes transcription, the possible implication of PMEPA1 in EMT and cell plasticity regulation is worth to be studied.

Here, we have identified, for the first time, *PMEPA1* as a COX2/PGF_2α_ up-regulated gene through the induction of TGFβ and we have deciphered its role in ovarian cancer progression. We have found that PGF_2α_ induced *TGFB1* and *PMEPA1* and we provide new evidence of its important role in ovarian cancer progression. Moreover, our results indicate that PMEPA1 is a critical regulator of epithelial plasticity, conferring a growth advantage in ovarian cancer cells.

## RESULTS

### Induction of *TGFB1* expression by PGF_2α_

TGFβ is a crucial factor for ovarian homeostasis as well as for tumorigenesis (22). Indeed, our gene expression analysis of the TCGA TARGET GTEx patient cohort RNA sequencing data using the UCSC XENA cancer browser shows a 2-fold increase of TGFB1 mRNA levels in ovarian tumor samples, as compared to normal ovarian tissue (fig. 1A). PGF_2α_ has an important role in the female reproductive system and is known to be produced in the ovary, as is produced also in ovarian tumors (11). Interestingly, we found that PGF_2α_ significantly increased TGFB1 mRNA levels in ovarian serous adenocarcinoma SKOV3 cells (fig. 1B). PGF_2α_ signals through the PTGFR, activating calcium-calcineurin (Ca^2+^/CaN) -nuclear factor of activated T cells (NFAT) (33). Indeed, Ca^2+^ ionophore A23287 significantly increased TGFB1 mRNA levels in SKOV3 cells. TGFB1 mRNA levels correlated to the PTGS2 mRNA levels (Pearson’s r = 0.20, p < 0.0001, n = 369), as they did with the PTGFR mRNA levels as well (fig. 1C). Since we already found TGFB1 was induced in response to Ca^2+^ signaling, we checked the mRNA levels of the NFAT-coding genes, to find that NFATC2 is more than 2 fold up-regulated in ovarian tumor samples as compared to healthy tissue (p < 0.0001, n = 514) (fig. 1D). Moreover, NFATC2 and TGFB1 mRNA levels correlated strongly in the same patient cohort (fig. 1E), as did also NFATC1 and TGFB1 (Pearson’s rho = 0.41, p < 0.0001, n = 369). As expected, TGFB1 levels also correlated with the *RCAN1 levels* (a *bona-fide* NFAT transcriptional target) mRNA levels in SKOV3 cells (fig. 1F).

**Fig. 1.**
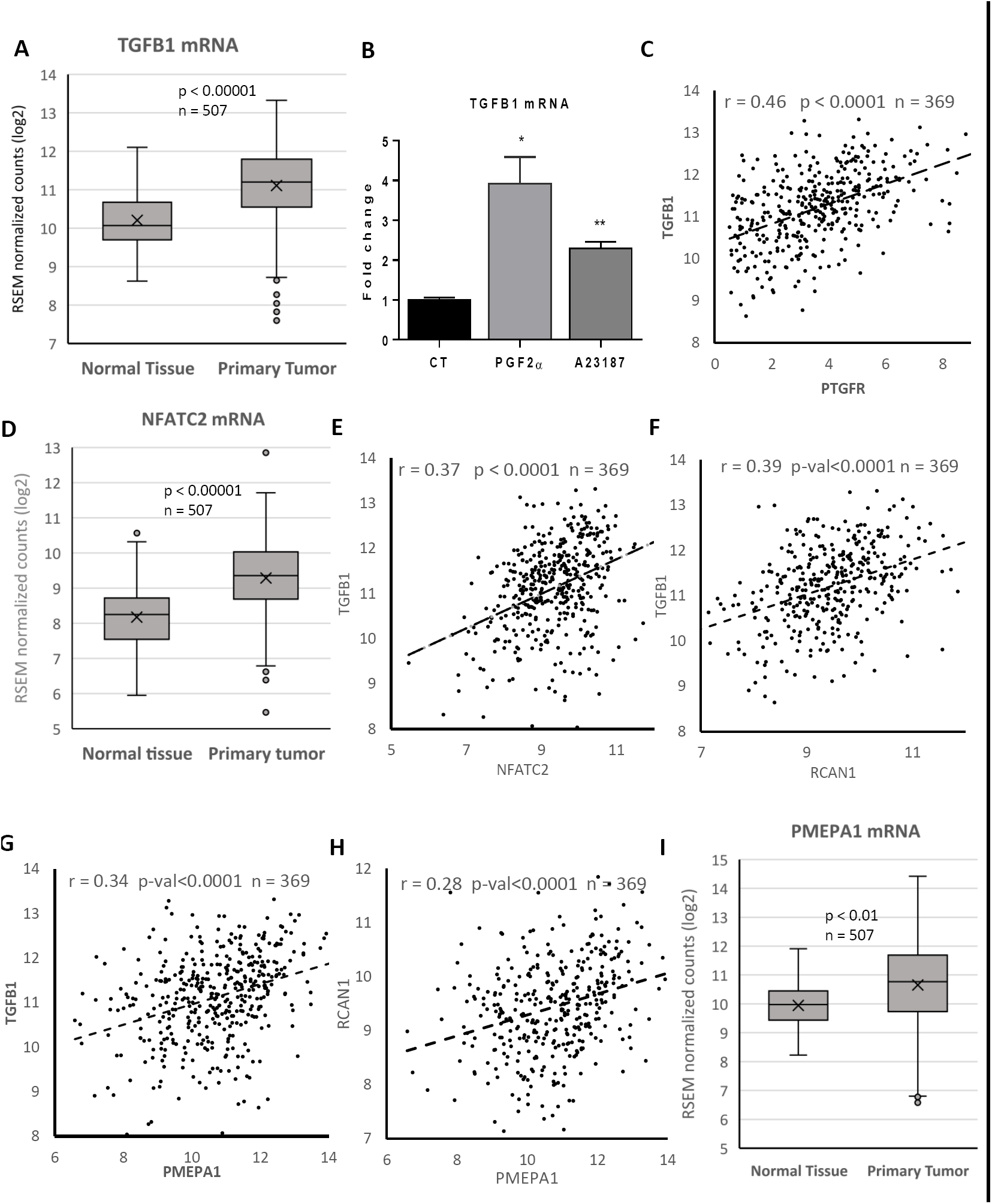
TGFβ is upregulated by PGF_2α_/NFAT in ovarian cancer. **A.** Normalized *TGFB1* mRNA levels in ovarian primary tumors and normal ovarian tissue, as calculated using the TCGA TARGET GTEx dataset in the UCSC Xena browser. Welch’s t-test, t = 9.398, p < 0.0001, n = 507. **B.** Relative TGFB1 mRNA levels estimated by RT-qPCR of control, 1μM PGF_2α_ or 1μM A23187 treated SKOV3 cells during 24h. *: p < 0.05 and **: p < 0.001. **C.** TGFB1 and PTGFR normalized mRNA levels, ovarian tumor and normal tissue samples XY dispersion from the same dataset as in A. The linear regression curve of the tumor samples values, Pearson’s correlation r and statistical significance are shown. **D.** Normalized NFAT mRNA levels, as in A. Welch’s t-test, t = 9.639, p < 0.0001, n = 507. **E.** XY dispersion of TGFB1 and NFATC2 mRNA levels as in C. **F.** XY dispersion of TGFB1 and RCAN1 mRNA levels as in C. **G.** XY dispersion of TGFB1 and PMEPA1 mRNA levels as in C. **H.** XY dispersion of PMEPA1 and RCAN1 mRNA levels as in C. **I.** Normalized PMEPA1 mRNA levels, as in A. Welch’s t-test: t = 6.464, p < 0.01, n = 507.

Along these lines, multiple NFATc1 binding sites were identified in the promoter region of TGFB1, using chromatin immunoprecipitation and mass sequencing (ChIP-seq) data of the Gene Transcription Regulation Database (GTRD), two of them in TGFB1 gene regulatory region located in its first intron (suppl. Fig. 1), Site IDs: 34541886-7. They were adjacent to c-fos binding sites, which could serve for co-activation, indicating a possible regulation of TGFB1 by PGF_2α_/NFAT, in agreement with the TGFB1 induction by PGF_2α_ or A23187 and expression correlation with RCAN1. Moreover, 3 EGR1-binding sites were found in the same regulatory region, site IDs 17860197-9. Remarkably, EGR1 is a transcription factor we found previously to be induced by PGF_2α_ (6). In accordance to this, we also observed that TGFB1 and EGR1 mRNA levels are correlated (r = 0.29, p < 0.0001, n = 369).

All these findings could indicate a close relationship between the COX-2/PGF_2α_/FP/Ca^2+^/NFAT pathway and TGFB1.

### PMEPA1 levels are elevated in patients’ ovarian tumor samples

PMEPA1 has been proposed as a TGFβ-induced gene able to convert TGFβ from a tumor suppressor to a tumor promoter (20). Indeed, we found that there is a strong correlation (Pearson’s r = 0.34) between TGFB1 and PMEPA1 mRNA levels in the TCGA ovarian cancer cohort (fig. 1G). A similar correlation was found between PMEPA1 and RCAN1 mRNA levels (fig. 1H), indicating the possible, direct or indirect, implication of NFAT in the PMEPA1 gene expression. Additionally, we found tumor PMEPA1 mRNA levels to be significantly higher than in normal ovary (fig. 1I).

To confirm these findings at the protein level, we performed immunohistochemistry analyses on normal tissue obtained from ovary fallopian tubes, fimbriae and peritoneal as well as in ovarian primary tumors and relapses. Primary ovarian tumors showed strong PMEPA1 staining in all of the tumor cells compared to normal tissues that showed diffuse expression in some epithelial cells (Fig. 2).

**Fig. 2.**
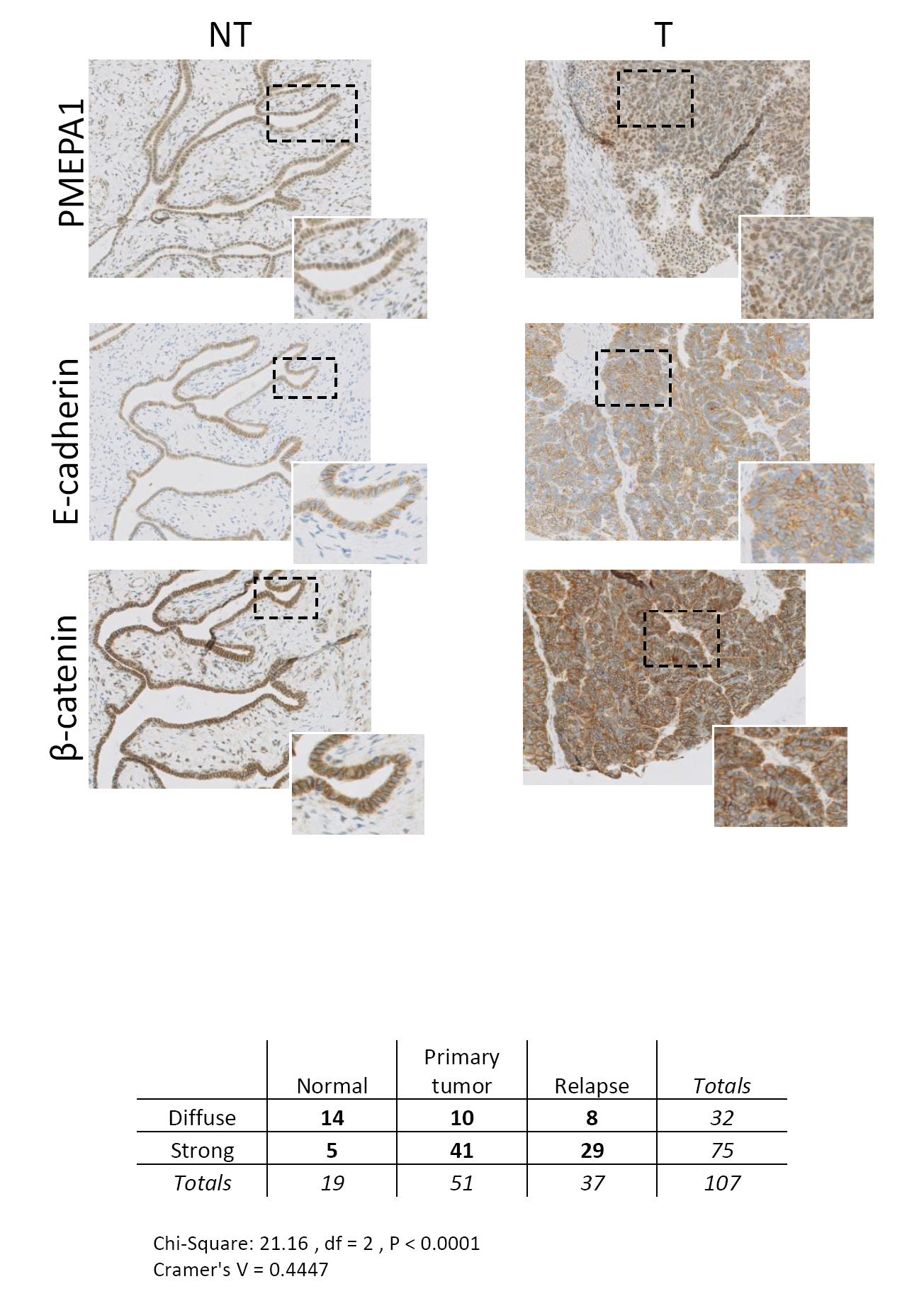
PMEPA1 expression is widely expressed in ovarian cancer. Representative images of PMEPA1 (top), E-cadherin (middle), and beta-catenin (down) expression by immunohistochemistry in normal **A**, and primary ovarian tumors **B**. Magnification 40×, inset 63×. The magnification areas are highlighted in the square. **C.** Quantitative analysis of PMEPA expression in normal, primary tumor and metastasis/relapse ovarian samples. The significance of the differences observed between the different groups was determined with a Chi-Square test P < 0.0001.

We decided to further explore the effect of PMEPA1 expression on patient survival. Since the patient cohort we used for PMEPA1 protein levels includes only high-grade tumors and most of them resulted positive for PMEPA1, no significant differences in patient survival according to PMEPA1 expression could be obtained. However, after analysis of the gene expression databases available (TCGA ovarian cancer), high PMEPA1 expression is associated with lower survival probability (suppl. Fig. 2). The above observations suggest PMEPA1 could have an important role in ovarian cancer progression and it could be considered as a potential biomarker for ovarian tumor characterization and patient stratification.

### Induction of *PMEPA1* expression by PGF_2α_

To study the PMEPA1 induction by TGFβ or PGF_2α_, we selected several ovarian cancer cell lines with different basal *PMEPA1* mRNA levels, from higher to lower, namely, SKOV3, OVCAR8, TOV112D and A2780 (fig. 3A), as quantified in the Cancer Cell Line Encyclopedia (34). Treatment with fluprostenol, a metabolically stable PGF_2α_ analog, was able to increase PMEPA1 mRNA levels, as did TGFβ treatment, in all the 4 cell lines tested. The combination of both treatments had a partially additive effect (Fig. 3B). We generated Skov3 derived cell lines transduced with lentiviral vectors able to express shRNAs that could knockdown *PTGFR* mRNA, thus decreasing PTGFR levels and signaling. When the resulting cell lines were treated with PGF_2α_ (fig. 3C) or fluprostenol (not shown) they failed to increase PMEPA1 mRNA levels as the scrambled shRNA carrying cells did, indicating that PMEPA1 induction by PGF_2α_ depended on the PG. Moreover, calcineurin/NFAT inhibition by cyclosporine A (CSA) could revert PGF_2α_- induced TGFB1 and PMEPA1 increase (fig. 3D). Indeed, CSA not only reverted significantly the two genes induction, it also reduced their levels in the absence of PGF_2α_, indicating a possible basal Ca2+/calcineurin signaling. Interestingly, the PTGFR induction of *PMEPA1*, but not *RCAN1*, a direct NFAT target, was reverted by co-treatment with an inhibitor of TGF-β type I receptor, LY2109761 (fig. 3E), indicating that basal TGFβR signaling may be necessary for basal and Ca^2+^ stimulated PMEPA1 expression.

**Fig. 3.**
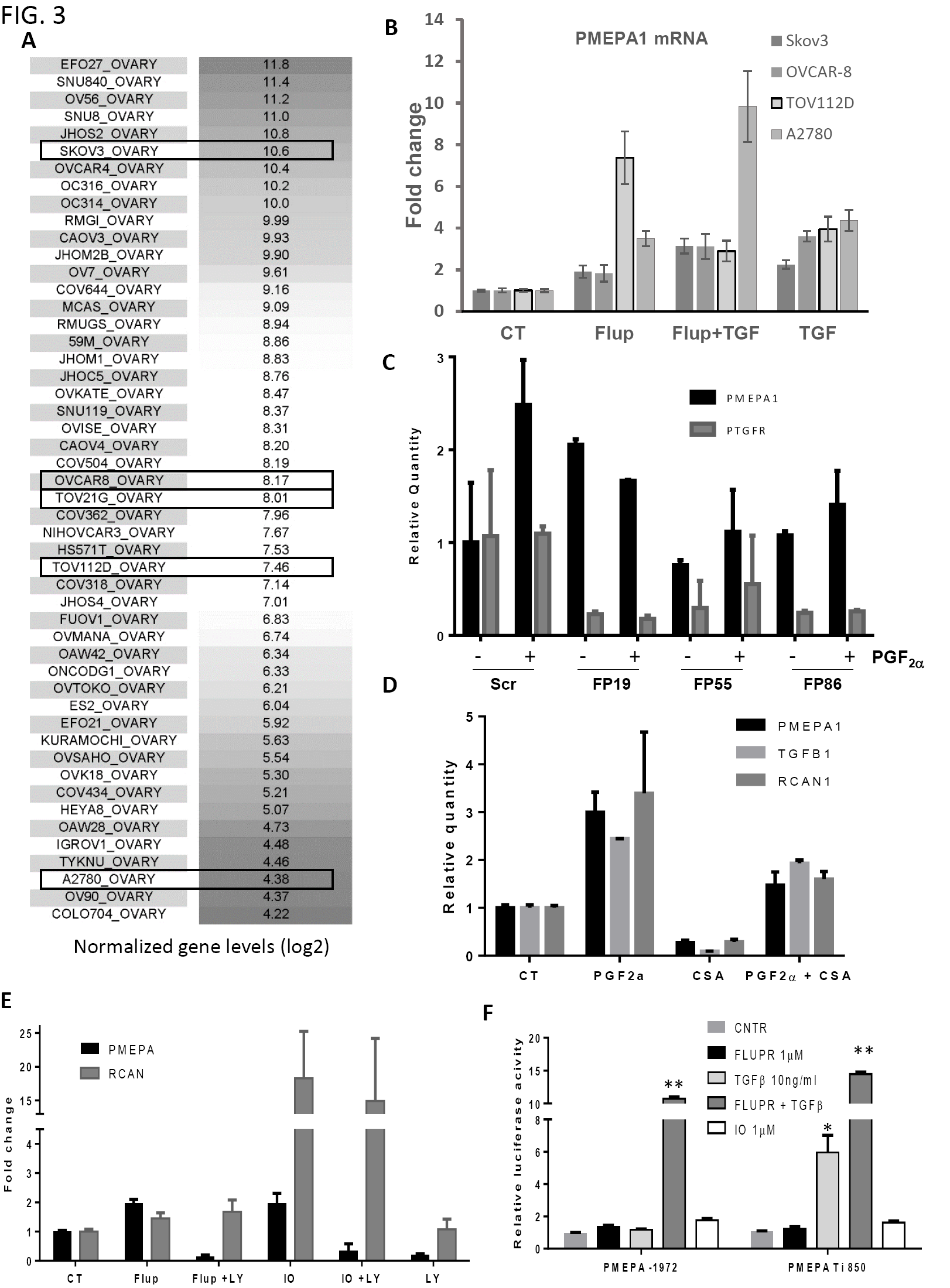
PMEPA1 is upregulated by cooperative action of TGFβ and PGF_2α_. **A.** Normalized PMEPA1 mRNA levels as calculated in Cancer Cell Line Encyclopedia (Broad Institute) in ovarian cancer cell lines ordered from highest to lowest expression. **B.** PMEPA1 mRNA levels quantification in the cells indicated after 24h of treatment with vehicle (CT), 1μM fluprostenol (flup), 5 ng/ml TGFβ, and the combination of the two (Flup+TGF). **C.** PMEPA1 and PTGFR mRNA levels quantification in SKOV3 derived cell lines stably expressing a scrambled shRNA (scr) or three PTGFR specific shRNAs (FP19, FP55, FP86) after 24h of treatment with vehicle (CT) or 1μM PGF_2α_. Only scr cells were able to significantly increase (p < 0.05) PMEPA1 mRNA with PGF_2α_ treatment. **D.** PMEPA1, TGFB1 and RCAN1 mRNA levels quantification in SKOV3 cells after 24h of treatment with vehicle (CT), 1μM PGF_2α_, 100ng/ml cyclosporine A (CSA) or combination of the two. CSA significantly reduced the levels of the three genes below the control levels (p < 0.05). PGF_2α_ significantly increased the three genes’ levels, while the combination of the two reduced the levels of the three genes, although in the case of TGFB1, the levels were significantly different from CT as well as from PGF_2α_. **E.** PMEPA1 and RCAN1 mRNA levels quantification in SKOV3 cells after 24h of treatment with vehicle (CT), 1μM Flup, 1mM A23187 Ca^2+^ ionophore (IO), 10μM LY2109761 (LY) or combinations. Both genes are induced by Flup or IO (p < 0.05). LY treatment only affects PMEPA1 levels reducing them below CT levels, even in combined treatments (p < 0.05). **F.** Luciferase reporter assays with SKOV3 cells transfected with the mentioned reporter constructs of the PMEPA1 promoter, and treated as shown for 24. TGFβ treatment only activated the PMEPA Ti 850 reporter (p < 0.05), while the combination of Flup and TGFβ activated both reporters (p < 0.001). IO or Flup alone did not have any activating effect.

To further investigate the role of Ca^2+^/NFAT and its possible cooperation with TGFβ signaling, we analyzed *PMEPA1* promoter activity. Two luciferase reporter constructs of the *PMEPA1* gene were used: the *PMEPA1* promoter fragment −1972*PMEPA1*-luc and PMEPA1 first intron pGL3ti-850 (24). The activity of pGL3ti-850 has been demonstrated to be potentiated by TGF-β as well as by the β-catenin/TCF4 complex (24,25). In SKOV3 cells, *pGL3*ti-850 was also stimulated by TGFβ treatment but not by fluprostenol alone, although it showed a strong synergistic activity with TGFβ. On the other hand, −1972*PMEPA1-luc* was only activated by the combination of fluprostenol and TGFβ (Fig. 3F). These results would indicate that SMADs and NFAT co-operatively stimulate PMEPA1 mRNA expression, while the PGF_2α_ induction of *PMEPA1* could be partially due to *TGFB1* induction.

We searched the GTRD ChIP-seq data and identified multiple NFATc1 binding sites, Site IDs 24982307-11, in the promoter and in the first intron regulatory regions of *PMEPA1*. Interestingly, we found several SMAD-2, SMAD-3 binding sites adjacent to the NFATc1 ones, as well as c-Jun and c-Fos (AP1, a common partner of NFAT) binding sites, which could offer co-activator binding to enhance transcriptional activation (suppl. figure 3).

All the above suggests that there could be a co-operation of NFAT with SMADs for basal levels maintenance as well as super-induction of *PMEPA1*. These results suggest that there is a positive feedback loop between PGF_2α_/NFAT and TGFβ for PMEPA1 expression. TGFβ induced *PMEPA1* expression can be further potentiated by the PGF_2α_/Ca^2+^/CaN pathway, both through its main promoter as by the first intron enhancer.

### PMEPA1 overexpression in cancer cells enhances cell growth

To investigate the biological effects of PMEPA1 in tumor cells, we generated ovarian cancer cell lines (Skov3, OVCAR8, A2780, TOV112D and TOV21G), stably overexpressing PMEPA1 and compared them with control cells carrying the empty vector (EV). Additionally, 5 knockdown SKOV3-Luc cell lines for PMEPA1 were generated. Knockdown or overexpression were confirmed by RT-qPCR, WB and immunocytochemistry of thin layer preparations (suppl. figure 4).

Interestingly, PMEPA1 overexpressing SKOV3 cells had a higher proliferative rate than SKOV3-EV cells (fig. 4A). Similar results were obtained in PMEPA1 overexpressing A2780, TOV112D (fig. 4B). PCNA protein levels increased accordingly, in agreement to this increased proliferation speed (fig. 5B). However, no difference in growth was observed in OVCAR8 cells (not shown). In contrast, proliferation rates of PMEPA1 knockdown cells were much lower compared to scrambled (SCR) control cells (fig. 4A). Anchorage independent survival revealed that PMEPA overexpressing cells survived better and formed bigger clusters that grew faster than EV cells (fig. 4C and D).

**Fig. 4.**
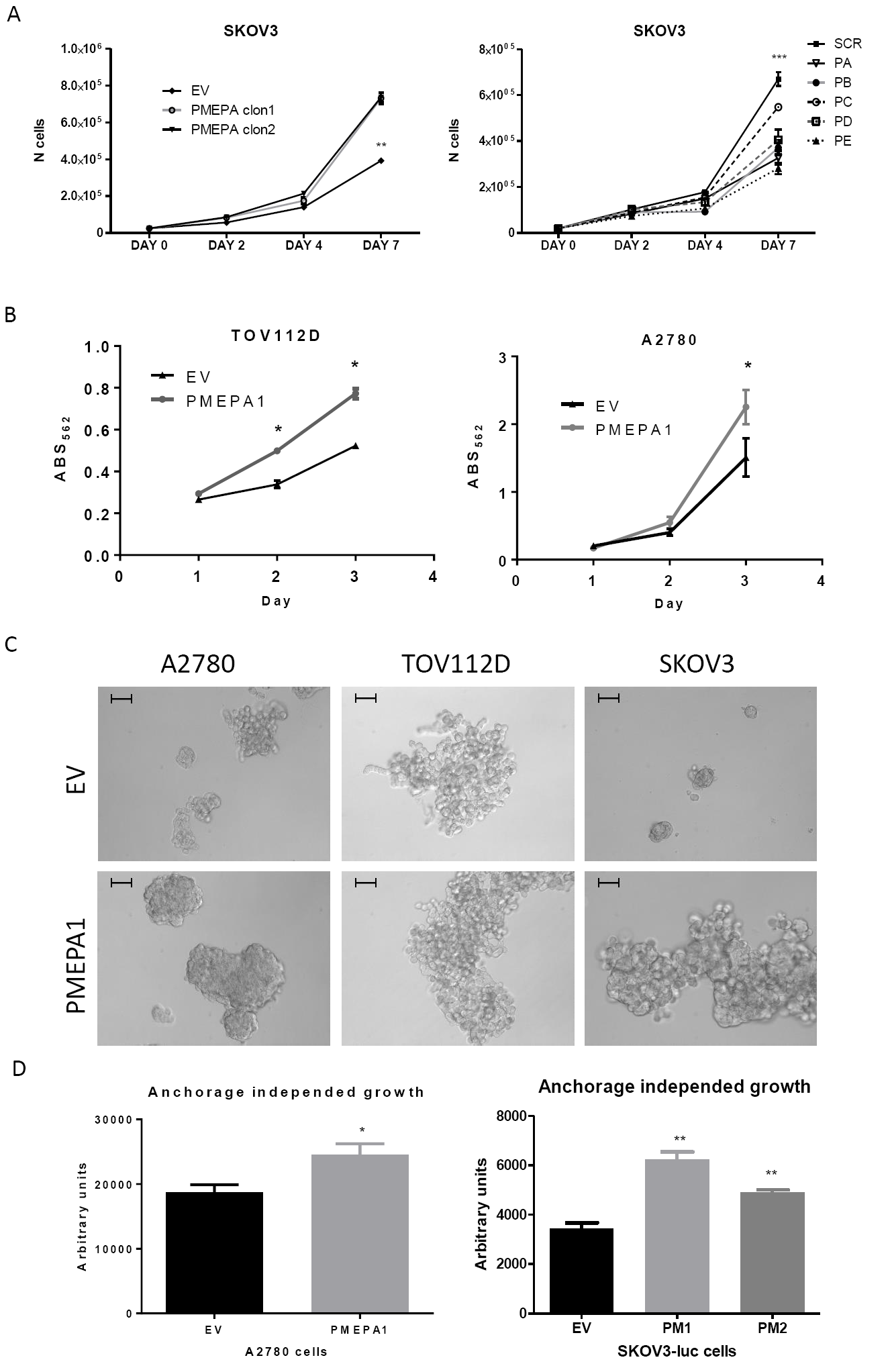
PMEPA1 overexpression in ovarian cancer cells enhances cell proliferation. **A.** Cell proliferation of 2 PMEPA1 overexpressing clonal SKOVE3 cell lines (*left panel*) and 5 knockdown (right panel) SKOV3 cell lines was estimated by cell counting using a hemocytometer. Average ± SEM are shown, n = 5. Growth curves were significantly different, p = 0.025 for overexpressing vs control and P < 0.001 for knockdown vs SCR. These results were also confirmed by crystal violet stain quantification of SKOV3 (not shown), A2780 and TOV112D EV and PMEPA cells (**B**). **C.** Anchorage independent growth assays were performed with PMEPA1 overexpressing and control A2780, TOV112D and SKOV3 cells that were monitored by light microscopy. Representative photos of the cells at 72 h after seeding are shown. Bars: 50μM. **D.** Cell survival and proliferation in the assays in (**C**) was estimated at 96 h after plating by Alamar blue assay. Stain reduction quantification is shown for A2780 and Skov3Luc (2 independent PMEPA1 overexpressing clones, PM1 and PM2). No significant differences were found for TOV112D (not shown). Average ± SEM are shown, n = 3.

**Fig. 5.**
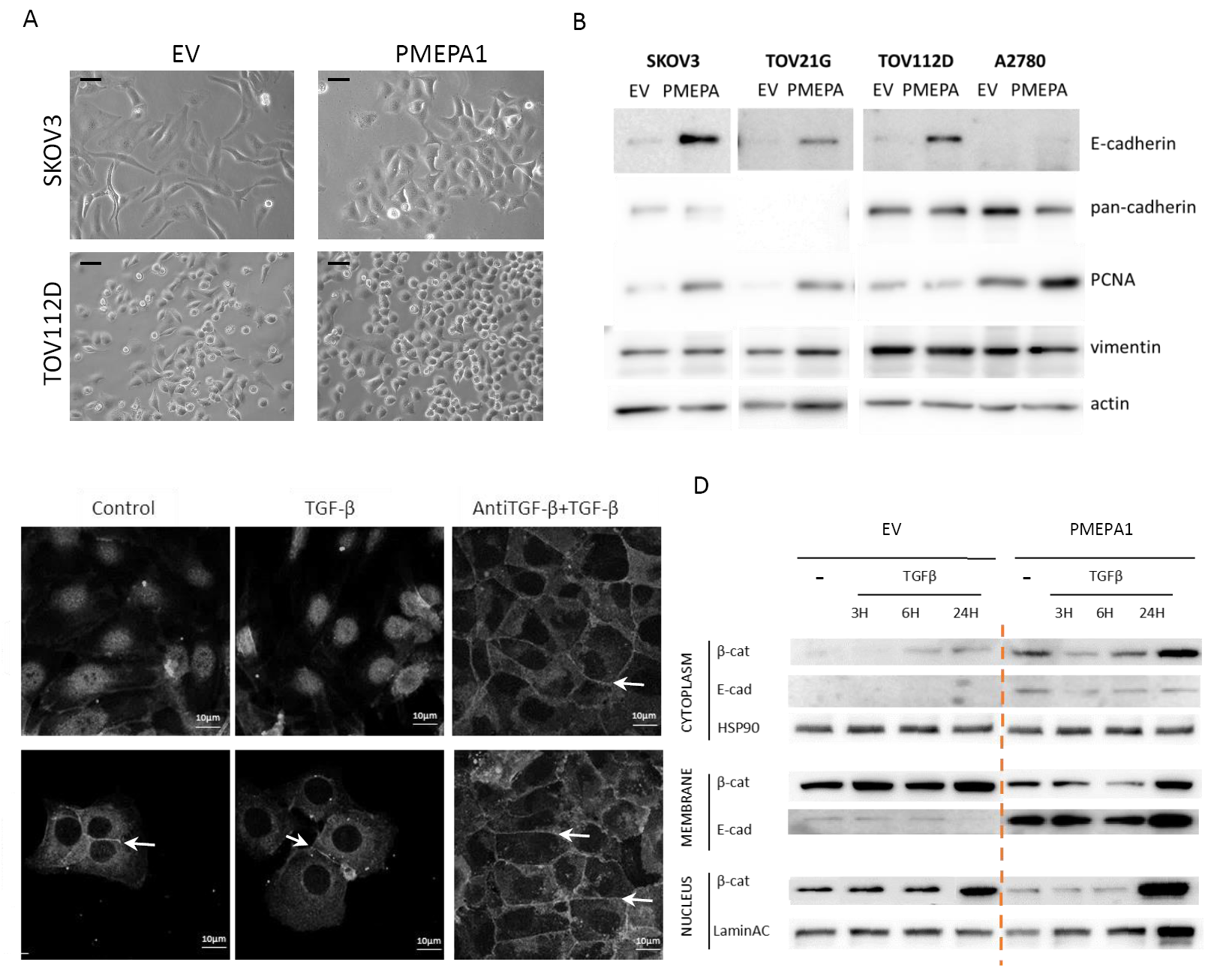
PMEPA1 overexpression favors a partially epithelial phenotype. A. Light microscopy images of SKOV3 and TOV112D -EV and –PMEPA1 cells in culture. Bars: 50μM. **B.** WB for the indicated proteins in SKOV3, A2780, TOV112D and TOV21G -PMEPA1 and -EV cells. **C.** Immunofluorescence for β-catenin of SKOV3-PMEPA1 and SKOV3-EV (EV) cells. Cells were treated with 5ng/ml of TGF-β or combination of 10μg/ml of recombinant Latency Associated Peptide (LAP-TGFβ) and 5 ng/ml of TGFβ for 24 hours. Arrows indicate β-catenin localization in cell-cell contacts. **D.** WB to detect the indicated proteins in nuclear, membrane and cytoplasmic extracts of the SKOV3-lucD6 EV and SKOV3-PMEPA1 cells.

### PMEPA1 overexpression favors a partially epithelial phenotype

Next, we addressed the cellular and molecular characteristics that may contribute to these important differences in ovarian tumor cells. SKOV3 cells have been classified as “intermediate mesenchymal phenotype cells”, through phenotypic and genomic attributes (35), in agreement with the fibroblast-like morphology the cells have in culture. On the other hand, OVCAR8 cells are classified as epithelial, while TOV112D and A2789 are classified as mesenchymal. Once SKOV3 cells were transduced with the PMEPA1 expression vector, morphological changes began to be noticeable towards a more epithelial phenotype (Fig. 5A). We observed similar changes in other ovarian cells lines like A2780, TOV21G and TOV112D upon transfection with the PMEPA1 construct while we did not find any changes in transduced Ovcar8 cells. As we detected increased cell-cell contacts, we checked E-cadherin protein expression that was higher in PMEPA1 overexpressing cells (Fig. 5B). This is in agreement with our finding that most ovarian tumors tested express both PMEPA1 and E-cadherin (fig. 2) and supported by the fact that *CDH1* is 40-fold more expressed in ovarian tumors as compared to normal ovarian tissue (suppl. fig. 5).

Taking into account the phenotypic change and E-cadherin increase, we decided to study β-catenin protein localization in PMEPA1 overexpressing cells, due to the known interaction between the adhesion molecule and the WNT signaling pathway member leading to the retention of the second in the cell membrane proximity (28). β-catenin localized both in the cytoplasm and nucleus of SKOV3-EV cells, while in PMEPA1 overexpressing cells it had a perinuclear localization as well as in cell-cell contacts, where E-cadherin can be found (fig. 5C, D). As expected, TGF-β treatment induced translocation of β-catenin to the nucleus in SKOV3 cells, while this effect was absent when PMEPA1 was overexpressed (Fig. 5C, D). Both the mesenchymal phenotype and β-catenin dissociation from the membrane could be due to autocrine or paracrine TGFβ signaling, since TGF-β blocking antibody (fig. 5C) or the LY2109761 TGF-β-RI inhibitor (not shown) induced similar changes as PMEPA1 overexpression, causing β- catenin membrane localization and changing to an epithelial-like morphology. Surprisingly, E-cadherin expression was not affected by TGF-β treatment in PMEPA1 cells (Fig. 5D). These results could indicate that PMEPA1 overexpression can revert downregulation of E-cadherin in ovarian cancer cells and decrease nuclear β-catenin.

The relationship between PMEPA1, E-cadherin and β-catenin, was supported by analysis of the above-mentioned patient tumor samples, showing that the samples that express PMEPA1, also express E-cadherin and β-catenin. β-catenin was in most cases detected in the plasma membrane and not nuclear (fig. 2). Taken together, these results support *in vitro* experiments, indicating that PMEPA1 high expression coincides with E-cadherin expression and blocks β-catenin nuclear translocation in ovary tumors.

### PMEPA1 overexpression affects the TGFβ signaling pathway

Increased phosphorylation of SMAD2/3 were observed (Fig. 6A) in PMEPA1 overexpressing cells. After TGFβ treatment, SMAD phosphorylation levels increased both in EV and in PMEPA1 overexpressing cells, but the fold increase was less pronounced in the case of SKOV3-PMEPA1 cells. On the contrary, we observed high levels of SMAD2/3 and P-SMAD2/3 proteins in SKOV3-shPMEPA1 compared to SCR (Fig. 6B). Interestingly, reduced levels of SMAD2/3 in nuclear extracts were found in SKOV3-PMEPA1 cells, while the membrane bound SMAD2/3 increased (fig. 6C). Besides, we tested if PMEPA1 affected SMAD-dependent transcription of TGFβ-target genes using the CAGA-Luc reporter in SKOV3 cells. Transcriptional activity was not significantly affected by PMEPA1 overexpression although we observed a tendency towards lower luciferase activity, both in control and in TGFβ-treated cells (Fig. 6D). In agreement to this, *SERPINE1* (PAI-1) and *EDN1* mRNA levels were lower in PMEPA1 overexpressing than in control cells (fig. 6E). On the other hand, PMEPA1 knockdown led to a significant reduction of the SMAD2/3-dependent reporter activity in untreated SKOV3 cells (fig. 6D) while no significant differences in SERPINE1 and EDN1 mRNA levels were found between the PMEPA1 knockdown and SCR cells. Interestingly, PMEPA1 mRNA expression significantly (p < 0.001) correlates with *SERPINE1, EDN1* and *TGFB1* (Pearson’s rho 0.47, 0.27 and 0.35, respectively, n = 427), as calculated with the UCSC Xena Cancer browser for TCGA Ovarian cancer. These results indicate that PMEPA1 is intrinsically related with the TGFβ pathway and should be considered as a necessary player for the pathway’s functions in cancer.

**Fig. 6.**
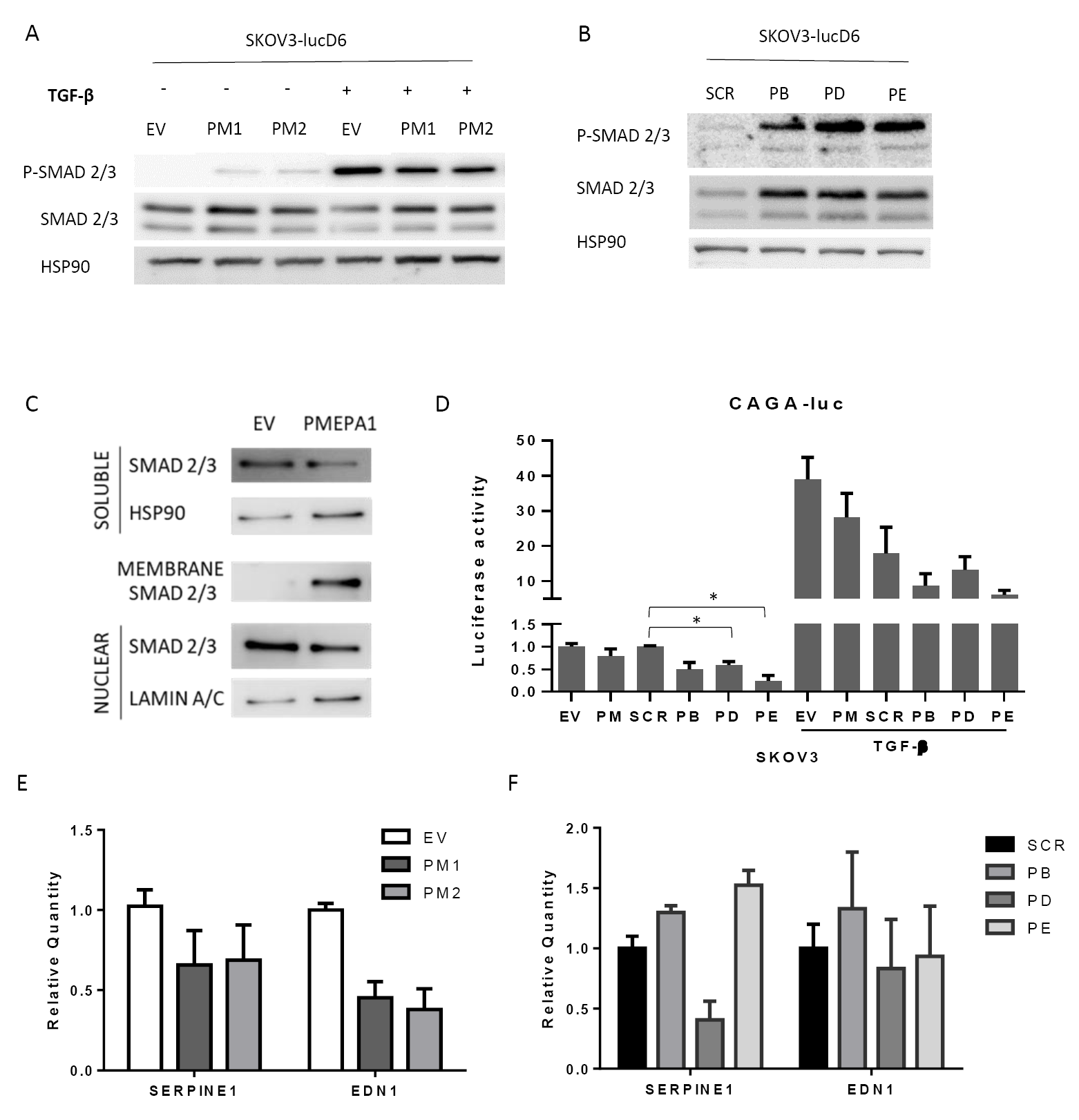
PMEPA1 overexpression affects the TGFβ signaling pathway. SMAD2/3 phosphorylation state as detected by WB in overexpressing (**A**) and knockdown (**B**) SKOV3 cells and their appropriated controls treated or not with 5ng/ml TGFβ for 1h. **C.** WB for SMAD2/3 protein levels in SKOV3-EV and -PMEPA1 cells nuclear, membrane and soluble extracts. **D.** Relative luciferase activity of ALK5-luc (CAGA reporter) transfected in PMEPA1, SKOV3 knockdown and their respective control cells, treated or not with 5ng/ml TGFβ during 6h. Relative SERPINE1 and EDN1 mRNA levels in PMEPA1 overexpressing (**E**) and knockdown (**F**) SKOV3 cells and their corresponding controls.

### PMEPA1 promotes tumor growth *in vivo*

Finally, to confirm that PMEPA1 promotes ovarian tumor growth, we tested PMEPA1 overexpressing or knockdown cell lines growth in mice (fig. 7). Several xenograft mouse models were used to address different aspects of ovarian cancer growth and metastasis. Nude mice were injected subcutaneously with 1×10^6^ SKOV3-Luc EV or PMEPA1 overexpressing cells and tumor growth was monitored over time. Tumor volume growth was much higher in SKOV3-PMEPA1 compared to SKOV-EV derived tumor that became palpable around day 55 (Fig. 7A). We were also able to follow and quantify tumor cell growth starting on the day of inoculation by bioluminescence. SKOV3-PMEPA1 xenografts showed a growth advantage when compared with SKOV3-EV xenografts from the beginning of the experiment (Fig. 7B). Mice were sacrificed and tumors from subcutaneous xenografts were extracted and photographed and we corroborated the mentioned results (suppl. fig. 6). SKOV3-EV derived tumors were smaller; in some cases, we were not able to isolate a tumor mass, although we could detect bioluminescent signal. To confirm these results, we also performed a similar experiment, with subcutaneous injection of TOV112D and A2780 control and PMEPA1 overexpressing cells, in R2G2 mice. In the case of TOV112D cells, PMEPA overexpression increased tumor initiation rates to at least 50%, comparing to 25% in EV cells, PMEPA1 tumors grew faster, reaching maximum allowed size at least a month earlier than the EV ones (suppl. Fig 7). On the other hand, only A2780-PMEPA1 cells, but not the EV, presented detectable growing tumors during a 15-week monitoring period (suppl. Fig 7).

**Fig. 7.**
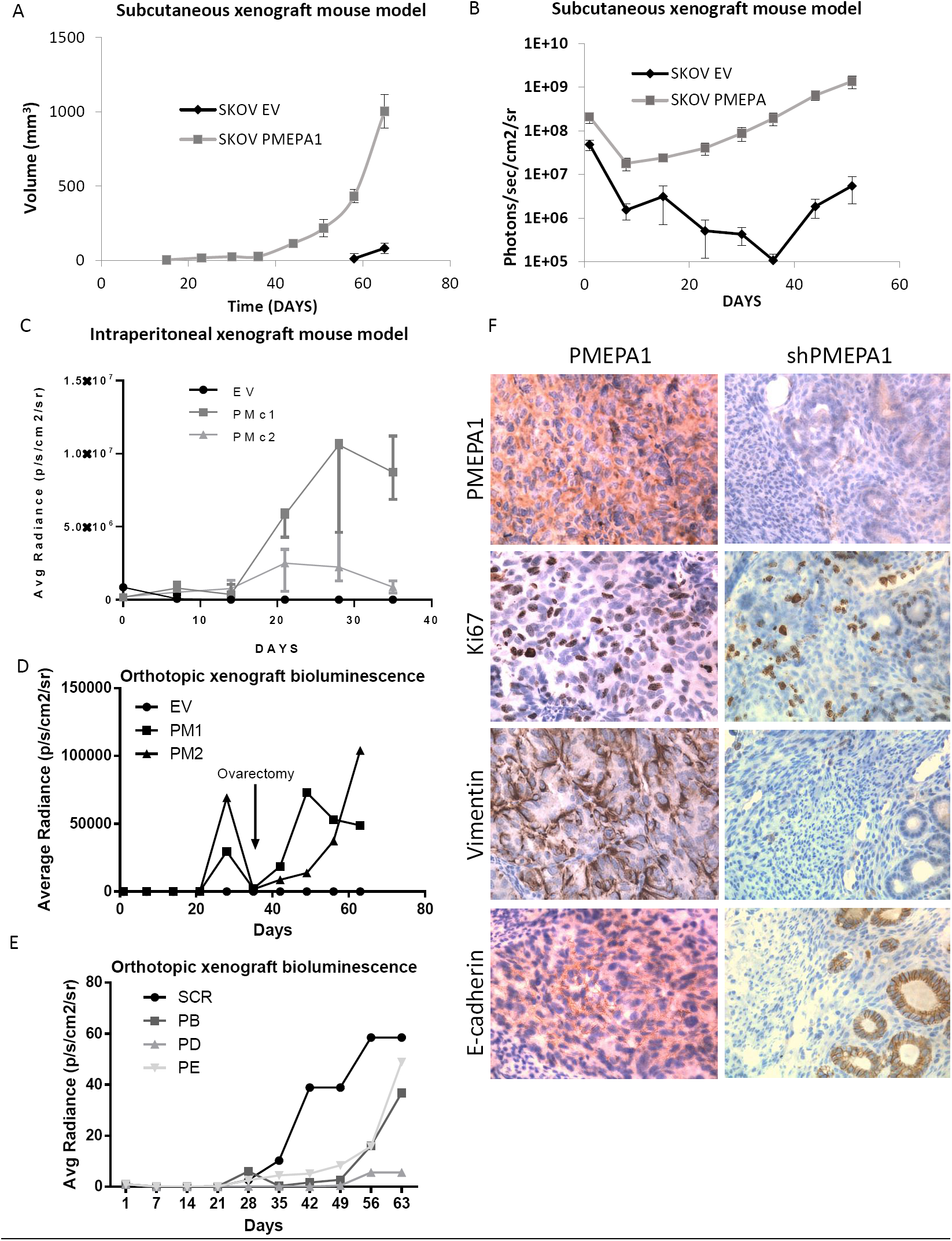
PMEPA1 promotes tumor growth *in vivo*. A. Subcutaneous xenograft Skov3Luc-EV and - PMEPA1 tumor volume, measured once a week, using a digital caliper. EV derived tumors became palpable around the 8^th^ week post inoculation. **B.** Bioluminescence Imaging (BLI) quantification of the xenografts in (A). **C.** Growth of intraperitoneal injected cells was measure by bioluminescence quantification with IVIS Lumina Imaging system. Orthotopic tumor xenografts growth estimation through bioluminescence quantification of PMEPA1 overexpressing (**D**) and knockdown (**E**) SKOV3Luc derived cell lines. Tumors were removed at 35 dpi, but BLI monitorization continued until day 63 in both experiments. **F.** IHC of tumors derived from *PMEPA1* overexpressing or knockdown SKOV3 cells, stained for the indicated proteins. Magnification ×400. B,C,D: Results are represented as logarithmic scale of Photons/sec/cm^2^/sr means ± SEM.

To further investigate the effects of PMEPA1 on tumor cell peritoneal dissemination, we injected intraperitoneally two SKOV3-PMEPA1 cell lines and SKOV3-EV cells. SKOV3-PMEPA1 xenografts had an important proliferative advantage, even more remarkable than in the subcutaneous experiments (Fig. 7C). To search for internal organ invasion, we extracted some organs (stomach, intestine, spleen, kidneys and pancreas) and peritoneal membrane. In mice that harbored SKOV3-PMEPA1, bio-luminescent signal could be detected in several internal organs, while in SKOV3-EV mice, all organs were negative (suppl. Fig. 8). Besides, peritoneal walls of SKOV3-PMEPA1 xenografts show localized luminescence signals corresponding to established metastases (suppl. Fig. 8). These results suggest that intraperitoneal dissemination and metastasis is facilitated by PMEPA1 overexpression.

Finally, to recapitulate all the steps of ovarian cancer growth, dissemination and metastasis, cells were injected orthotopically into the ovary. SKOV3-PMEPA1 xenografts showed a strong growth advantage compared with SKOV3-EV, which could hardly form intra-ovary tumors (Fig. 7D). More importantly, PMEPA1 knockdown SKOV3 grew even less, reaching even lower levels of BLI signal than SCR and much less than SKOV3-PMEPA1 xenografts (Fig. 3E) indicating that even the low PMEPA1 levels of control cells are important for tumor growth. Thus, the difference between overexpressing and silencing PMEPA1 in SKOV3 cells resulted in differences of about 20,000 fold in bioluminescence intensity.

Finally, to investigate the potential for these cells to develop metastasis, primary tumors were removed on day 35 (red arrow) and mice were monitored for bioluminescence, indicating metastatic tumor growth. A week after the operation, bioluminescence could be detected again, indicating cells had already metastasized (Fig. 7E). Thus, PMEPA1 gives a clear advantage to ovarian tumor cells to adapt and grow in all the tested conditions. These in vivo results and the PMEPA1 expression in patient high-grade tumors strongly suggest that PMEPA1 could be an accurate biomarker of prognosis in ovarian cancer.

## DISCUSSION

Elevated expression of Cyclooxygenase 2 (COX2) has been associated with development several cancers (36) and it is well established that COX2 inhibition can help prevent cancer (3). Besides, the importance of Calcineurin/NFAT signaling pathway in *COX2* transcriptional induction and its involvement in colon tumor properties were already demonstrated by our group (1,6). PGF_2α_ on the other hand, has only recently been associated with tumor progression in colorectal (6,37) and endometrial cancer (38,39). Since it is naturally produced in the ovary and capable to produce strong effects on the epithelium (40) it would be logical to assume that it may have an important role also in ovarian cancer. Remarkably, although PTGFR levels are lower in ovarian tumors than in normal ovary, ovarian cancer patients with higher PTGFR mRNA levels exhibit lower survival probabilities than patients with lower PTGFR levels (suppl. Fig 9). These findings support studies on the implication of PTGFR in ovarian and other cancer types, since it appears to be an interesting pharmacological target against ovarian cancer dissemination and metastasis.

The TGFβ superfamily plays an important role in ovarian function and pathogenesis (41) while, remarkably, mutations in genes of this pathway are infrequent in ovarian cancer (27). Importantly, we observed *TGFB1* is up-regulated in ovarian cancer. We are the first to describe the induction of TGFB1 by PGF_2α_ in cancer cells and our observation is supported by one earlier publication observing this induction in the bovine *corpus luteum* (42). Not surprisingly, we found PTGFR and TGFB1 levels to be strongly correlated in ovarian tumors.

Furthermore, there is a strong correlation between the expression of TGFB1 and all the components of the COX2-PGF_2α_-PTGFR-NFAT pathway tested. Moreover, we reported for the first time the association of Ca^2+^/Calcineurin/NFAT signaling with the transcriptional control of *TGFB1*. Furthermore, a cooperation of NFAT and EGR1 in the TGFB1 upregulation is highly plausible, since we have found several EGR1-binding sites in the TGFB1 promoter region, adjacent to NFAT-binding sites and we have already shown that EGR1 is also induced by PGF_2α_ to regulate gene expression (6).

We thus offer data that, for the first time, show the convergence between two pathways of great importance in cancer progression, TGFβ and COX2. COX2 products, such as PGF_2α_, are able to induce TGFB1 transcription. This could further contribute to explain many of the effects of COX2 and PGs in the tumor setting, such as EMT (43), metastasis (44) and immune evasion (45) and could point out a potential interest in develop combined therapies including TGFβ and COX2 modulators in ovarian cancer.

As mentioned above, ovarian tumors rarely acquire mutations in the TGFβ pathway, although this cytokine is up-regulated in tumors over normal tissue. Thus, we sought to find a gene product that could allow tumor growth, involved in the evasion of the TGFβ tumor suppressor effects. One that fits this premise would be PMEPA1, already proposed to be a “molecular switch that converts TGF-β, normally a tumor suppressor, to a tumor promoter” (20).

Indeed, we found that PMEPA1 is elevated in most ovarian tumors, its expression correlating with that of TGFB1. Remarkably, we found that PMEPA1 is induced also by PGF_2α_/NFAT axis, as TGFB1, reaching a synergistic effect between the two pathways. Our data on the pro-tumoral effect of PMEPA1 in ovarian cancer are also supported by the fact that high PMEPA1 mRNA levels are associated with lower survival rate of patients. PMEPA1 overexpression increased tumor cell growth both *in vitro* and *in vivo*, as we demonstrate using different cell lines and *in vivo* tumor models, while it’s knockdown had the opposite effect. Thus, silencing PMEPA1 resulted in a much reduced tumor growth *in vivo*. More interestingly, our results also suggest that intraperitoneal dissemination and metastasis is facilitated by PMEPA1 overexpression. Given the fact that intracellular TGFβ signaling moderately decreased, we believe that PMEPA1 expressing cells are able to affect the tumor stroma without suffering the negative effects TGFβ could induce.

PMEPA1 overexpression altered cell morphology, prompting a more epithelial phenotype, although TGFβ production by the cells increased. This altered morphology could be due to the up-regulation of E-cadherin we observed. Indeed, PMEPA1 was able to elevate functional E-cadherin levels and thus concentrate part of the β-catenin in the intercellular junctions, while other epithelial markers failed to be induced. It has been already proposed that E-cadherin expression can have a positive effect on tumor aggressiveness and metastasis (46), which would perfectly agree with our results. Moreover, we observed that PMEPA overexpressing cells, not only had elevated growth capacity on substrate, but also anchorage independent growth, in accordance with previous reports attributing this role to E-cadherin (47). E-cadherin could be degraded upon different stimuli, as TGFβ(48), and thus lose association with β-catenin, that could translocate to the nucleus (49). SKOV3-EV cells show β-catenin nuclear localization probably due to basal autocrine TGF-β signaling (50) and absence of membrane E-cadherin. Even β-catenin remains in a cytoplasmic localization in PMEPA1 overexpressing cells, as we are the first to demonstrate. Indeed, after TGF-β treatment, more β-catenin translocated to the nucleus in SKOV3-EV cells but it remained cytoplasmic in SKOV3-PMEPA1 cells, indicating that PMEPA1 overexpression can block the β-catenin nuclear translocation. A plausible explanation to this phenomenon could be the fact that β-catenin can depend on SMAD3 to translocate to the nucleus (51,52). This is also supported by the fact that we found the same epithelial morphology and β-catenin cytoplasmic localization when we treated cells with a latency associated peptide, inhibiting TGFβ or a TGFβR inhibitor. Indeed, we also found that the overexpression of PMEPA1 reduced nuclear SMAD2/3, as already described in other cell systems (19,20,23).

It is well established that E-cadherin expression and decreased cell mobility are common epithelial cell characteristics, while upregulation of N-cadherin, vimentin and zinc-finger domain proteins (SNAI1/SNAIL, SNAI2/SLUG), among others, are often linked to a mesenchymal-like phenotype (53). A remarkable case is the ovarian surface epithelial (OSE) cells, in which overexpression of E-cadherin induces a number of epithelial characteristics and markers associated with malignant transformation and tumor progression (54). Remarkably, both primary and metastatic ovarian carcinomas express E-cadherin, in contrast to normal ovarian surface epithelium, which rarely expresses E-cadherin (55,56). Further work should be done to elucidate the mechanisms through which PMEPA1 upregulates or avoids downregulation of E-cadherin and if these depend exclusively on TGFβR signaling. Interestingly, PMEPA1 effects on growth can be observed only in ovarian carcinoma cell lines with mesenchymal or intermediate mesenchymal phenotype but not in OVCAR8 cells that are classified as epithelial. This supports idea that the effects of PMEPA1 in favoring an epithelial phenotype are linked to those on growth advantages.

A growing amount of evidence shows that PMEPA1 has a pro-tumoral role in different tissues (18,57) while its pro-metastatic role is more controversial (58). However, our findings show that PMEPA1 gives a clear pro-metastatic advantage to ovarian cancer cells. We believe that the PMEPA1 positive or negative effect on tumor growth could be related to a TGF-β/β- catenin or E-cadherin dependency of each tissue. Using a high grade ovarian tumor patient cohort, we found strong PMEPA1 expression in most tumors, perfectly correlating with E-cadherin and β-catenin high expression. Bigger cohorts, including lower grade primary tumor biopsies should be used to confirm if PMEPA1 is linked to metastasis and its prognostic value. All the above could indicate that PMEPA1 can tweak TGFβ signaling while making tumor cells produce more, upregulate E-cadherin protein levels and reduce β-catenin nuclear localization, thus increasing cell plasticity and proliferation.

In conclusion, we show here a novel interaction between two signaling pathways important in cancer, COX2/prostaglandins/NFAT and TGFβ that leads to tumor progression and metastasis in ovarian cancer. Moreover, we demonstrate the importance of PGF2a and PMEPA1 in these processes. Further investigation on the role of PMEPA1 in ovarian cancer is necessary, since it might be an appropriate target for therapy and an accurate biomarker for prognosis and patient stratification, useful in clinical practice.

## MATERIALS AND METHODS

### Ovarian samples

A series of 19 normal, 51 primary tumors and 37 metastatic/relapse ovarian samples were collected at the MD Anderson Cancer Center Biobank (Madrid). The sample characterization was performed by a pathologist (ARS), who determined the histological cancer subtype according to the World Health Organization (WHO) criteria (59), and the stage and grade (supplemental table 1). The local ethical committee of the MD Anderson Cancer Center approved the study, and a complete written informed consent was obtained from all patients.

### Cell lines

SKOV3-lucD6 cells, stably expressing Firefly Luciferase, were obtained from Caliper Life Sciences. SKOV3, TOV112D and TOV21G cells were obtained from ATCC and A2780 cell line was provided by Sigma-Aldrich. Human embryonic kidney HEK293-FT cells were purchased from Invitrogen. OVCAR8 cell line was a generous gift from Dr. JM Cuezva (Centro de Biología Molecular Severo Ochoa). All cell lines were grown in the recommended conditions and cell line purity was verified with the StemElite ID system (Promega).

### Reagents

All generic reagents were from Invitrogen or Sigma-Aldrich. Oligonucleotide and antibody details can be found in supplementary tables 2 and 3.

### Plasmids

pLenti CMV/TO Hygro (w214-1) was a gift from Eric Campeau & Paul Kaufman (Addgene plasmid # 17484). For PMEPA1-pLentiCMV/TO construction, the primers 5’- ATGGATCCATGCACCGCTTGATGGGG-3’ and 5’-TACCTCTAGACTAGAGAGGGTGTCCTTTC-3’, carrying XbaI and BamHI restriction sites were used for PCR amplification of the PMEPA1 open reading frame (transcript 001, human PMEPA1 cDNA, NM_199170 Origene). <u>pALK5/SMAD3-luc</u> (CAGA-luc): consisted of 12 tandem repeats of the upstream SMAD3-binding element from human PAI-1 promoter linked to a viral minimal promoter and to a luciferase gene (60). The *PMEPA1* promoter fragment −1972*PMEPA1*-luc (from +67 to – 1972) and PMEPA1 first intron pGL3ti-850 (from +447 to +1294) reporter constructs were a generous gift from Dr. S. Itoh (Showa Pharmaceutical University, Tokyo, Japan) (24). <u>shPMEPA1 scrambled shRNA (scr) and shPTGFR in pLKO.1-puro:</u> human PMEPA1 and PTGFR MISSION shRNA Bacterial Glycerol Stock (SHCLNG-NM_020182 and SHCLNG-NM_000959 respectively) from SIGMA-Aldrich shown in supplementary table 4. <u>TCF4-WT</u>: TCF7L2_pLX307 was a gift from William Hahn & Sefi Rosenbluh (Addgene plasmid # 98373).

#### Lentiviral vector transduction

Lentiviral plasmids co-transfection and lentivirus collection was done according to Addgene protocols. All lentiviral plasmids and the packing plasmids (pCMV 8.2 and pMDG) were co-transfected into HEK293FT cells using Lipofectamine 2000 (Invitrogen). Lentiviral particles were collected 72 h after transfection and used to transduce target cells. 48 hours after transduction cells were trypsinized and selected using Hygromycin (200 μg/ml) (for the pLenti system) or Puromycin (2 μg/ml) (for the pLKO.1 puro shRNA system).

**Cell proliferation assays** were performed plating 20000 cells in p35 wells. Cells were plated in 3 or 4 different wells and each day of the experiment cells from a different well were counted. Cell proliferation was also quantified by Crystal Violet staining. Cells were washed two times with PBS before they were fixed in 5% Glutaraldehyde in PBS for 20 minutes. Fixing solution is removed and cells were stained with 0,5% Crystal Violet in water and 50% methanol for 20 minutes. Crystal Violet solution was washed by immersion in water tap 3 times and cells are left to dry. Crystal Violet stain was dissolved in 10% Acetic Acid and optical density was measured at 570nm.

**Substrate independent cell growth assays** were performed by seeding 50,000 cells/well in ultra-low attachment surface 24-well plates (Costar) in normal growth medium. Cells were monitored for 96 h by light microscopy. At the end of the experiment relative cell numbers were estimated by Alamar Blue staining (Invitrogen).

#### RNA extraction and RT-qPCR

Total RNA was isolated using TRIzol reagent (Invitrogen) and following the manufacturer’s protocol. qPCR analysis was performed using the GoTaq 2-Step RT-PCR system (Promega) as described (6), using ABI PRISM 7900HT Fast qPCR system. Relative mRNA levels to the housekeeping *HPRT* (Hypoxanthine-guanine phosphoribosyltransferase) gene and to the experimental control point were calculated using the 2^−ΔΔCT^ formula from the values obtained: (where Ct are threshold cycles, ΔCt = Ct_gene_ – Ct_HPRT_ and ΔΔCt = ΔCt_problem_- ΔCt_control_).

**Immunofluorescence** was performed as described before (6). Images were taken using a Zeiss Axioscop2 plus with a color CCD camera or the Zeiss LSM710 confocal microscope.

**Western blotting** was performed as described before (6). For nuclear/cytoplasmic/membrane fractionation, medium was removed from the culture dishes, cells were washed with cold PBS and lysed for 15 min on ice with buffer A (10mM HEPES (pH 7,6), 10mM KCl, 0,1mM EDTA, 0,1mM EGTA, PhosSTOP and COMPLETE phosphatase and protease inhibitors, Roche), hypotonic buffer that extracts soluble proteins only. Cells were scraped, gently vortexed and lysates centrifuged to separate supernatant (cytoplasmic fraction) and pellet. The pellet washed and then resuspended in buffer A and NP-40 was added to a final concentration of 0.5%. Lysates were gently vortexed and centrifuged to separate the membrane fraction (supernatant) from the nuclear (pellet). The pellet was resuspended, washed and lysed with Buffer C (20mM HEPES (pH 7,6), 0,4mM NaCl, 1mM EDTA, 1mM EGTA, PhosSTOP and COMPLETE). Protein quantification was performed by Bradford assay. Analysis of proteins was done as previously described.

#### Luciferase assays

Cells were grown in p24 multiwell plates and were transfected with reporter plasmids and RenillaSV40 used for transfection efficiency normalization. Stimuli were added after 24 hours. Cells were lysed with Passive Lysis Buffer 5x from Dual-Luciferase Reporter Assay kit (Promega) and luciferase assay was performed according to the manufacturer’s protocol. Luciferase and Renilla luciferase activities were read in a P96 Nunclon multiwell plate (white surface) in FLUOstar OPTIMA equipment (BMG LABTECH). Luciferase activity was quantified obtaining the ratio samples/control.

#### Tumor growth in nude mice

In all cases, the minimum cell number indicated for each xenograft model was used. In the case of A2780 and TOV122D cell lines, 10^6^ and 5×10^4^ cells were injected subcutaneously in 6 week old female Rag2/Il2rg Double Knockout mice (R2G2, ENVIGO, France). For the SKOV3-Luc subcutaneous and intraperitoneal xenograft models, 6-week-old female Swiss Nude mice (Crl:NU (Ico)-Foxn1^nu^, Charles River Laboratories) were injected with 1×10^6^ cells, following the instructions of the provider of the cells, Caliper LS. Mice were weighted once a week and the tumor volume was estimated with a digital caliper by measuring: length × width × height. Tumor growth estimation was made by injecting the luciferase substrate D-Luciferin (150mg/kg of the mice body weight) intraperitoneally and bioluminescence was measured on anesthetized mice (1.5% isoflurane, Abbott) using a Xenogen IVIS^®^ Lumina CCD camera (Caliper Life Sciences). The luminescent signal was quantified with Living Image 3.2 software and expressed as photons/s (Average Radiance). In the orthotopic mouse model of ovarian cancer, animals were injected with 5×10^5^ in the left ovary. Tumor cell number estimation was made as mentioned above using the Xenogen IVIS^®^ Lumina CCD camera. The animal research described in this manuscript complies with Spanish Act (RD 53/2013) and European Union Legislation (2010/63/UE). The protocols were approved by the Animal Care Committee of the Centro de Biología Molecular Severo Ochoa (CBMSO, Madrid, Spain.

#### Histological analysis and immunohistochemistry

Tumors from mice and patients were fixed in 4% phosphate buffered formalin (pH 7.4) and 2 μm paraffin-embedded sections were immunostained, performed as previously described (6). Antibodies used for immunohistochemistry: anti-PMEPA1 (1:50) as primary antibody (Sigma-Aldrich) and secondary antibody conjugated with HRP (Envision + Dual link System HRP, Dako). Finally, sections were developed using DAB solution (Liquid DAB + substrate chromogen system, DAKO K3468) and images were taken with a LEICA DMD108 Digital Microimaging Device (Leica Microsystems).

### Databases and genomics analysis

Patient tumor and normal samples gene expression analysis was performed through the UCSC Xena http://xena.ucsc.edu using public access data from The Cancer Genome Atlas (TCGA, http://cancergenome.nih.gov/), the National Cancer Institute’s TARGET Data Matrix (https://ocg.cancer.gov/programs/target) and the Genotype-Tissue Expression (GTEx) project(61), to compose the TCGA TARGET GTEx cohort.

ChIP-seq data and metadata were analyzed using the Gene Transcription Regulation Database (GTRD, http://gtrd.biouml.org/) (62). The regulatory (promoter) region of each gene was estimated using the UCSC Genome browser (http://genome.ucsc.edu) CpG Islands, H3K27Ac marks and DNAse hypersentitivity regions visualization tools. Graphic output of both databases’ queries showing the gene-of-interest region was composed for the images in the supplementary figures 1 and 3.

### Statistical analysis

Results are expressed as mean ± SEM. The Student’s t test, the ANOVA test or Welch’s t-test were used for comparisons, where necessary. *p<0,05 and **p<0.01 denote statistical significance. For gene expression correlation/covariance, Pearson’s correlation coefficient was calculated. The statistical analysis was performed using the GraphPad Prism 4.0 statistical software.

## Acknowledgments

Tissue samples were obtained with the support of MD Anderson Foundation Biobank (record number B.0000745, ISCIII National Biobank Record). We appreciate Maria Chorro and Marta Ramiro for technical assistance. We appreciate assistance of the CBMSO Confocal Microscopy Service and Animal facility Service personnel.

